# A pathogenic missense mutation in kainate receptors elevates dendritic excitability and synaptic integration through dysregulation of SK channels

**DOI:** 10.1101/2023.06.21.545929

**Authors:** Toshihiro Nomura, Sakiko Taniguchi, Yi-Zhi Wang, Nai-Hsing Yeh, Anika P Wilen, Charlotte C.M. Castillon, Kendall M. Foote, Jian Xu, John N. Armstrong, Jeffrey N. Savas, Geoffrey T. Swanson, Anis Contractor

## Abstract

Numerous rare *de novo* variants that cause neurodevelopmental disorders (NDDs) occur within genes encoding synaptic proteins, including ionotropic glutamate receptors (iGluRs). However, in many cases it remains unclear how damaging missense variants affect brain function. Here we determined the physiological consequences of an NDD causing missense mutation in the *GRIK2* kainate receptor (KAR) gene, that results in a single amino acid change p.Ala657Thr in the GluK2 receptor subunit. We engineered the equivalent mutation in the mouse *Grik2* gene, yielding a GluK2(A657T) mouse, to better understand the human disorder and determine how hippocampal neuronal function is disrupted. Synaptic KAR currents in hippocampal CA3 pyramidal neurons from heterozygous A657T mice exhibited slow decay kinetics, consistent with incorporation of the mutant subunit into functional receptors. Unexpectedly, CA3 neurons demonstrated elevated action potential spiking due to down-regulation of the small conductance Ca^2+^ activated K^+^ channel (SK), which mediates the post-spike afterhyperpolarization (AHP). The reduction in SK activity resulted in increased CA3 dendritic excitability, increased EPSP-spike coupling and lowered the threshold for the induction of LTP of the associational commissural (AC) synapses in the distal dendrites of CA3 neurons. Pharmacological inhibition of SK channels in wild-type (WT) mice increased dendritic excitability and EPSP-spike coupling, mimicking the phenotype in A657T mice and suggesting a causative role for attenuated SK activity in aberrant excitability observed in the mutant mice. These findings demonstrate that a disease-associated missense mutation in *GRIK2* leads to altered signaling through neuronal KARs, pleiotropic effects on neuronal and dendritic excitability, and implicate these processes in neuropathology in patients with genetic NDDs.

## Introduction

Rare *de novo* variants causative for neurodevelopmental disorders (NDDs) occur in a variety of genes that code for synaptic proteins including ionotropic glutamate receptors (iGluRs)^1^. iGluRs are central to excitatory neurotransmission and critical for normal development of the brain. *De novo* pathogenic variants have been identified in genes that encode each of the four classes of iGluRs and gain and loss of function variants cause disorders with a spectrum of clinical features that include intellectual disability and global developmental delay ^2-13^. The number of damaging iGluR gene mutations identified in patients with NDDs continues to grow rapidly, but it remains to be determined how specific variants cause changes to brain development or function that result in cognitive, motor, and other phenotypes.

In the *GRIK2* KAR gene, damaging missense variants alter channel function, protein expression, and trafficking of assembled complexes to the plasma membrane ^6,7^. The first such variant, p.Ala657Thr, was discovered in a child whose primary clinical features were global developmental delay, intellectual disability, and an ataxic gait ^7^. More recently, additional individuals with similar neurodevelopmental symptoms were discovered to have the same variant ^6^. The *GRIK2* p.Ala657Thr variant occurs at a position in the GluK2 receptor protein critical to function that is part of a highly conserved set of residues, the SYTANLA<u>A</u>F motif, which form the distal segment of the M3 transmembrane pore-forming domain and affects gating of the receptor-channel. An analogous mutation in the mouse *Grid2* iGluR gene was the first such reported *de novo* variant and is causative for an ataxic phenotype in the “Lurcher” strain of inbred mice. This mutation in *Grid2* results in a constitutively active GluD2 delta subunit that causes Purkinje neuron degeneration^14,15^, and has also been identified in humans ^16^. Pathogenic variants at the analogous alanine in *GRIA* AMPA receptor genes result in NDDs that include autism spectrum disorder (ASD) in some children ^9,11-13^.

KARs are expressed across a wide range of brain regions in both excitatory and inhibitory neurons but in most are excluded from postsynaptic densities and therefore do not mediate fast, phasic synaptic depolarizations. Genetic and pharmacological studies implicate KARs in the fine-tuning of neurotransmitter release from presynaptic terminals ^17-19^. Other unconventional activities include promoting the functional expression of K-Cl cotransporter isoform 2 (KCC2) ^20^ and actively suppressing the post-spike afterhyperpolarization (AHP), which elevates neuronal excitability ^21-23^. KARs also modulate important synaptic properties, neuronal morphology, maturation and plasticity during development ^24^. While the mechanisms that KARs play in modulating circuit development are not yet fully known, and may be quite heterogeneous, in many cases these involve non-canonical, G protein-mediated signaling pathways distinct from ionotropic function of the receptor ^22,25,26^ and tonic, non-phasic activation by ambient glutamate ^27^. Therefore, damaging KAR gene variants have the potential to disrupt key aspects of neuronal signaling and development of the CNS, consistent with clinical phenotypes in children with *GRIK2* mutations ^6,7^.

In this study, we have examined the consequences of aberrant KAR signaling in a mouse line harboring the mutant *Grik2* allele that generates the p.Ala657Thr alteration, referred to as GluK2(A657T) mice. Our examination was focused on determining how receptors containing the mutant subunit disrupt excitability and synaptic function in hippocampal CA3 pyramidal neurons, where the signaling properties of KARs have been extensively characterized ^28-30^. Surprisingly, spontaneous action potential (AP) frequency was elevated in CA3 neurons. There was also a decrease in the AHP in CA3 neurons of GluK2(A657T) mice that correlated with enhanced dendritic excitability and increased AP coupling of distal associational-commissural (AC) synapses. Elevated dendritic excitability also boosted the spread of backpropagating APs (bAPs) and AC synapses exhibited a reduced threshold for the induction on NMDA receptor dependent long-term potentiation (LTP). Altered dendritic excitability was due to an apparent functional downregulation of small conductance Ca^2+^ activated K^+^ channels (SK) in A657T mice. These results demonstrate that a *Grik2* mutation that models a human genetic NDD causes profound disruptions in hippocampal function that at least in part arise from processes that alter dendritic excitability.

## Results

### Synaptic KARs are modified in GluK2(A657T) mice

To understand how the Ala657Thr mutation affects brain function, we engineered a mouse with a single amino acid swap from alanine (A) to threonine (T) in the GluK2 receptor subunit using CRISPR editing. Heterozygous GluK2(A657T) mice were viable but no living homozygous mice were observed, suggesting that a biallelic Ala657Thr mutation in *Grik2* is embryonic lethal. In hippocampal sections GluK2/3 immunoreactivity appeared similar to that in wild-type (WT) mice in the dentate gyrus (DG) and CA3 regions where KARs are abundantly localized (Figure 1A). Western blot analysis of the P2 and P3 fractions (enriched for large mossy fiber (MF) boutons^31^) from hippocampal extracts demonstrated that there were no major disruptions in synaptic expression of GluK2/3 protein (P2 fraction: n = 3, p = 0.24; P3 fraction: n = 3, p = 0.54; *t*-test) (Figure 1B & C). As a discovery-based screen of potential differentially expressed proteins in GluK2(A657T) mice, we used Tandem Mass Tags (TMT)-based quantitative mass spectrometry (MS) analysis to compare the CA3 proteomes in WT and A657T heterozygous mice (Figure 1D & E). We found that protein levels were highly similar between the two genotypes, notably, only ∼0.89% of CA3 proteins were significantly altered in the A657T bulk CA3 extracts. Bioinformatic Gene Ontology enrichment analysis of the significantly elevated or reduced proteins revealed no significantly enriched terms. Importantly, NMDA and AMPA receptor proteins were not different in WT and A657T mice. These data demonstrate that introduction of the *Grik2* p.Ala657Thr allele has only a minor effect on the CA3 proteome and that synaptic proteins are expressed at the same levels as in the WT littermates.

**Figure 1:**
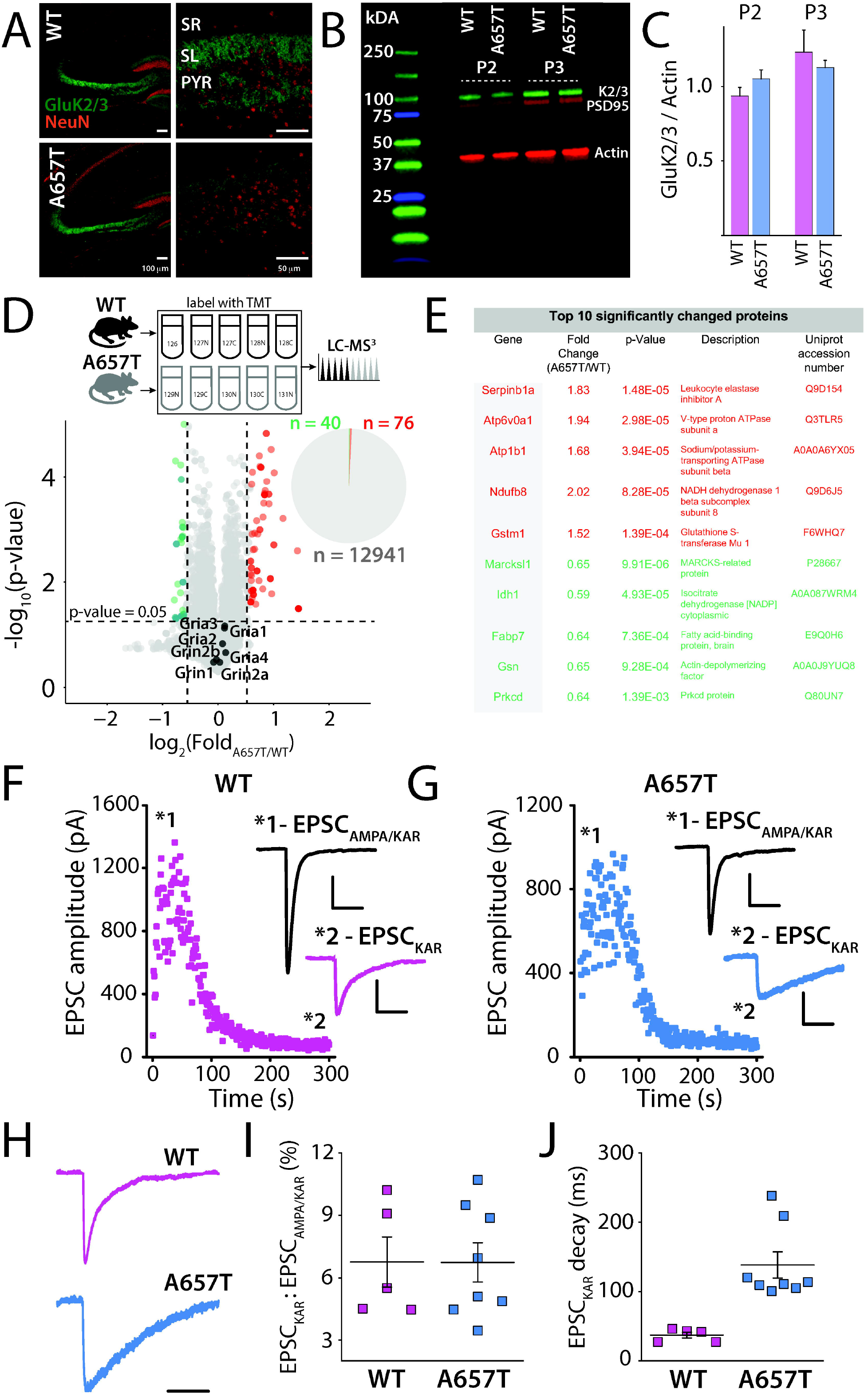
Synaptic kainate receptor mediated currents and analysis of the CA3 proteome in A657T mice. (**A**) Immunohistochemical analysis of KAR expression in the hippocampus. GluK2/3 labeling in the CA3 region was abundant in the termination zones of the mossy fiber axons in the stratum lucidum (SL) in both WT and A657T mice. SR denoted the stratum radiatum and PYR the pyramidal cell layer of CA3. Calibration, left panels: 100 µm; right panels: 50 µm. (**B**) Representative Western blot of GluK2/3, PSD95 and Actin in the P2 and P3 fractions from hippocampus of WT and A657T regions. (**C**) Quantification of Western blots (**D**) Top panel: Experimental design to compare WT and A657T CA3 proteomes. Bottom Panel: Volcano plot depicting comparison of relative protein abundance in WT and A657T mutant CA3. The major NMDAR and AMPAR protein subunit isoforms are indicated on the plot. Pie chart showing only ∼0.89% CA3 proteins are differentially expressed in A657T CA3. n = 5 mice per genotype. Significance was assessed using Student’s t-test. (**E**) Table of top 10 significantly differentially expressed proteins (red, upregulated; green, downregulated (**F**) Mossy fiber EPSC recorded in CA3 neurons stimulating at 1Hz to elevate release and measuring EPSC before and after the application of AMPA receptor selective antagonist GYKI53655 (50μM) in WT and (**G**) A657T mice. Inset shows the EPSC_AMPA/KAR_ prior to antagonist and the residual EPSC_KAR_ (magnified) after AMPA receptor blockade. Calibration EPSC_AMPA/KAR_: 250 pA, 50ms; EPSC_KAR_: 50 pA, 50 ms. (**H**) Deactivation kinetics of EPSC_KAR_ are slowed in A657T mice. Representative traces pf EPSC_KAR_ are shown. (**I**) Analysis of EPSC amplitudes normalized to the amplitude of the total mossy fiber EPSC. (**J**) Analysis of the decay of the EPSC_KAR_.

To test whether the mutant subunit assembled into functional synaptic KARs, we recorded KAR-mediated excitatory postsynaptic currents (EPSC_KAR_) from CA3 pyramidal neurons, which localize postsynaptic GluK2-containing receptors exclusively to MF synapses ^28,29^. Voltage clamp recordings were made from CA3 pyramidal neurons and mossy fiber inputs were electrically stimulated in the *stratum lucidum*. AMPA receptors were inhibited with GYKI-53655 (50 μM), leaving the isolated EPSC_KAR_ (Figure 1 F & G). The relative amplitude of EPSC_KAR_ normalized to the total EPSC_AMPAR/KAR_ was not different between WT and A657T mice, consistent with normal expression of GluK2-containing KARs in the mutant mice (WT: 6.8 ± 1.2 %, n = 5 cells, 3 mice; A657T: 6.7 ± 0.9 %, n = 8 cells, 5 mice; *p* = 1.0; Mann-Whitney) (Figure 1H & I). As predicted ^7^, the decay kinetics of EPSC_KAR_ were significantly slower in recordings from the A657T mice, demonstrating that mutant subunits are generated and incorporated into synaptic KARs (WT: 37.0 ± 4.1 ms, n = 5 cells, 3 mice; A657T: 138.7 ± 18.9 ms, n = 8 cells, 5 mice; *p* = 0.0016; Mann-Whitney) (Figure 1 J). Prior recordings of mutant GluK2A657T receptors in recombinant systems have demonstrated a constitutive steady state current in the nominal absence of agonist ^7^. To determine whether we could detect a standing current in CA3 neurons in A657T mice we measured the change in holding current after application of the non-specific AMPA/KAR antagonist NBQX (50 μM). We found that in recordings from CA3 neurons in WT mice there was no change in holding current after NBQX (WT: 2.74 ± 3.10 pA, n = 22 cells, 7 mice) whereas there was a small but significant current detected in A657T recordings (A657T: 16.6 ± 3.58 pA, n = 16 cells, 4 mice; *p* = 0.0012, Mann-Whitney)(Figure S1 A&B)

We also assessed whether parameters of mossy fiber synaptic physiology known to be disrupted in GluK2 null mice ^17^ were altered in the A657T mice. There were no changes apparent in paired pulse facilitation (Figure S1 C&D), short term plasticity during trains of stimuli at 1 Hz (Figure S1 E & F) or in NMDA receptor-independent MF long-term potentiation (LTP) compared to WT mice (Figure S1 G & H). These results demonstrate that mutant GluK2(A657T) subunits readily incorporate into neuronal KARs and slow deactivation kinetics of the EPSC_KAR,_ uncover a standing current in CA3 neuron recordings in the slice, but do not affect expression of synaptic receptors or their influence on presynaptic forms of potentiation.

### CA3 neurons are more excitable in A657T mice

To test if the A657T mutation affects the functional properties of neurons in the CA3 region, we first recorded spontaneous action potential firing (action currents) in CA3 neurons. The majority of neurons in WT mice were silent throughout the recording period (5 min), with only 40% (17/42) of cells firing at least one action potential. In contrast, 84% (27/32) of CA3 pyramidal neurons in A657T mice were spontaneously active, and the average spike frequency was higher than that in neurons from WT mice (WT: 0.19 ± 0.11 Hz, n = 42 cells, 17 mice; A657T: 0.38 ± 0.21 Hz, n = 32 cells, 8 mice; *p* = 0.000012; K-S) (Figure 2A & B).

**Figure 2:**
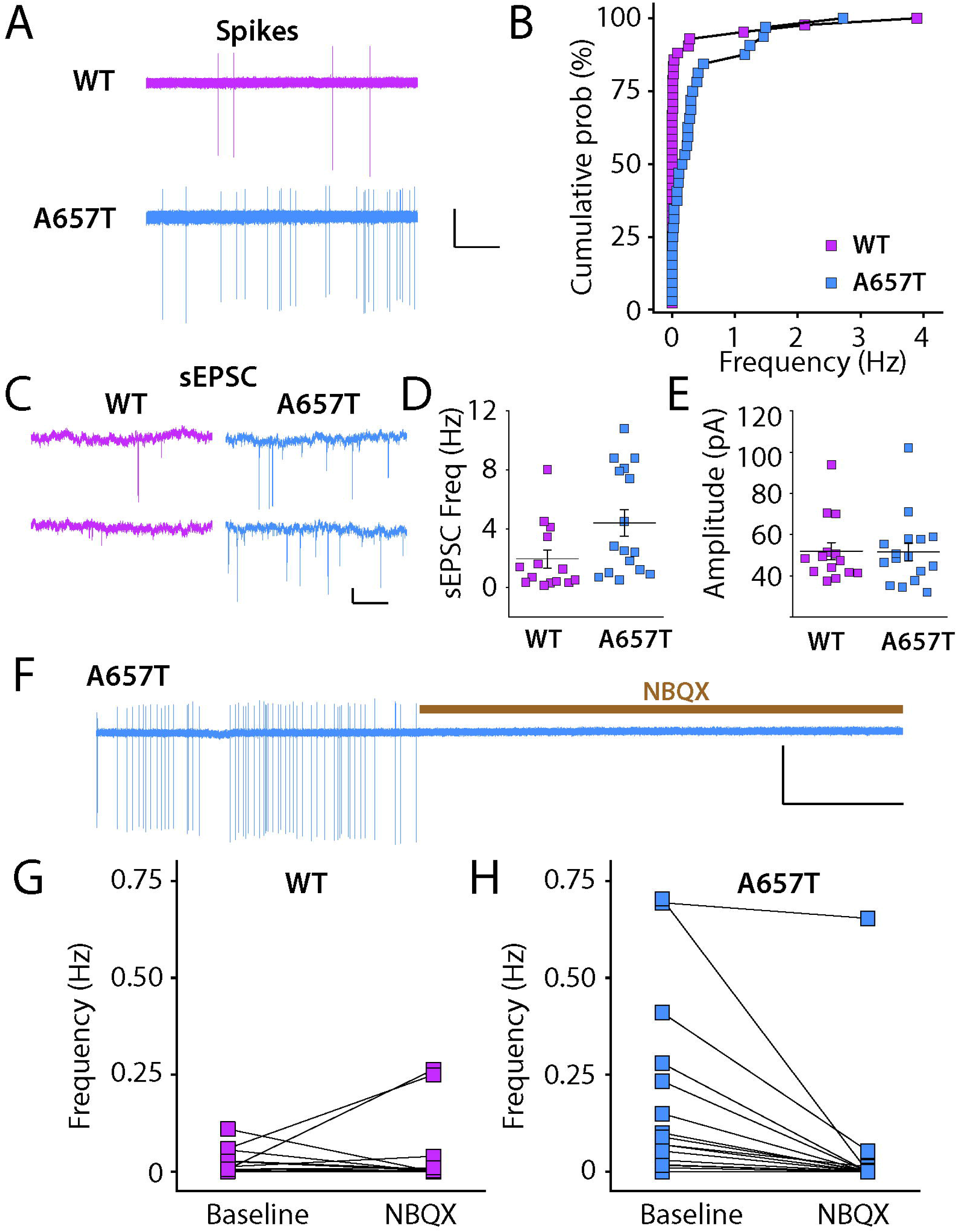
CA3 neurons are more excitable in A657T mice. (**A**) Representative action currents (spikes) recorded from CA3 neurons in cell-attached mode from WT and A657T mice. Calibration: 50 pA, 25 s (**B**) Cumulative distribution of spike frequency recorded in CA3 neuron from WT and A657T mice. (**C**) Example traces of sEPSCs recorded in CA3 neurons in WT and A657T mice. Calibration: 50pA, 1s (**D**) sEPSC frequency in CA3 is elevated in A657T mice (**E**) Amplitude of sEPSCs is not affected in A657T mice. (**F**) Example continuous trace of spikes recorded in CA3 neurons prior to and after glutamate receptor antagonist application NBQX, (50 µm) (**G**) Analysis of spontaneous action current frequencies before and after NBQX in WT and (**H**) A657T mice.

Spontaneous action potential firing is dependent on both the intrinsic properties of pyramidal neurons as well as integration of inhibitory and excitatory synaptic input ^32^, either of which could be affected in CA3 neurons in A657T mice. We assessed these parameters in recordings of synaptic currents and intrinsic properties of CA3 neurons. We determined the resting membrane potential (RMP) was more depolarized (WT: -68.0 ± 1.7 mV, n = 27 cells, 8 mice; A657T: -61.2 ± 1.5 mV, n = 24 cells, 7 mice; *p* = 0.0069; Mann-Whitney)(Figure S2 A) and the input resistance (R_in_) was lower (WT: 207.3 ± 10.9 MΩ, n = 28 cells, 14 mice; A657T: 168.7 ± 8.1 MΩ, n = 44 cells, 18 mice; *p* = 0.0035; Mann-Whitney) in CA3 pyramidal neurons from A657T mice compared to WT, which could be attributed in part to the standing KAR current we had measured in CA3 neurons. In contrast, there were no significant differences in rheobase (WT: 161.56 ± 26.7 pA, n = 16 cells, 6 mice; A657T: 166 ± 16.2 pA, n = 15 cells, 6 mice, *p* = 0.49; Mann-Whitney), or AP threshold (WT: -39.4 ± 2.0 mV, n = 16 cells, 6 mice; A657T: -43.1 ± 1.3 mV, n = 15 cells, 6 mice, *p* = 0.32; Mann-Whitney) in A657T mice. When APs were evoked by somatic depolarization, the input-output (I-O) AP curve was indistinguishable between WT and A657T mice (WT: n = 16 cells, 6 mice; A657T: n = 13 cells, 5 mice; two-way ANOVA, *F*_(1,27)_ = 0.0003, *p* = 0.99) (Figure S2 B & C). We also found that voltage-dependent conductances that contribute to the AP, including the total K^+^ current (I_K+total_) and the total Na^+^ (I_Na+total_), were not different in CA3 neurons of A657T mice (not shown). Together, we interpreted these data as evidence that intrinsic properties likely play a relatively minor role in the elevated AP firing observed in CA3 neurons in A657T mice.

We next tested if synaptic excitation and inhibition are altered by measuring spontaneous postsynaptic currents (sEPSCs and sIPSCs). The frequency of sEPSCs was significantly elevated in A657T mice (WT: 1.9 ± 0.6 Hz, n = 14 cells, 6 mice; A657T: 4.4 ± 0.9 Hz, n = 16 cells, 5 mice; *p* = 0.020; Mann-Whitney) without a change in sEPSC amplitude (*p* = 0.93; Mann-Whitney) (Figure 2C-E). In contrast, neither the frequency nor the amplitude of sIPSCs were different between CA3 pyramidal neurons in slices from WT and A657T mice (Figure S2 D-F). Elevated AP spiking of CA3 therefore alters the balance of synaptic excitation to synaptic inhibition in these neurons.

Altered integration of synaptic input could generate the elevated AP frequency observed in our initial experiments. To test this possibility, we recorded spontaneous action currents while blocking EPSCs with the AMPA/kainate receptor antagonist NBQX (50 µM). Spontaneous spike firing in CA3 neurons from A657T mice was mostly eliminated by inhibition of excitatory synaptic transmission with NBQX (Figure 2F & H) with little apparent effect in WT mice (Figure 2G). These data suggest that increased excitatory synaptic activity might contribute to spiking of CA3 neurons and increase recurrent network excitability in A657T mice.

We next sought to determine whether hippocampal neuron spiking might be elevated *in vivo* in A657T mice. We assessed the expression of the immediate early gene cFos as a proxy for neuronal activity. Acute bouts of voluntary wheel running are known to elevate cFos expression in all hippocampal subregions ^33-35^. We therefore tested whether cFos labeling is differentially elevated in mutant mice in CA3 pyramidal neurons and their upstream partners in the dentate gyrus (DG) after a single exposure to a running wheel in their homecage (Figure 3A). Mice in the A657T and WT groups did not demonstrate any differences in the total average running distance (WT: 0.809 ± 0.166 km, n = 13 mice; A657T: 0.768 ± 0.211 km, n = 6 mice, *p* = 0.77, Mann-Whitney) (Figure 3C). We analyzed cFos labeled cell density in the DG upper blade (DG_UB_) DG lower blade (DG_LB_) and in regions CA3a-c (Figure 3A-B). In each of these subregions the density of cFos labeled neurons was elevated in A657T mice (DG_UB_: 0.86 ± 0.15 x 10^-3^ /μm^2^, 1.86 ± 0.33 x 10^-3^ /μm^2^, p = 0.030; DG_LB_: 0.63 ± 0.14 x 10^-3^ /μm^2^, 1.47 ± 0.35 x 10^-3^ /μm^2^, p = 0.026; CA3c: 1.20 ± 0.24 x 10^-3^ /μm^2^, 2.73 ± 0.65 x 10^-3^ /μm^2^, p = 0.046; CA3b: 1.24 ± 0.20 x 10^-3^ /μm^2^, 3.13 ± 0.68 x 10^-3^ /μm^2^, p = 0.012; CA3a: 0.90 ± 0.20 x 10^-3^ /μm^2^, 3.26 ± 0.79 x 10^-3^ /μm^2^, p = 0.0047; WT: n = 13 mice, A657T: n = 6 mice, Mann-Whitney) (Figure 3B). Correlating the total density of cFos labeled neuron in the DG or CA3 with the total running distance also demonstrated a strong correlation between distance run by each mouse and density of cFos in each of these regions (Pearson’s r: DG, WT: 0.64 DG, A657T: 0.60; CA3, WT: 0.70; CA3, A657T: 0.70) (Figure 3D & E), highlighting that the animal’s activity was directly related to the expression of cFos in each genotype but was elevated in A657T mice.

**Figure 3:**
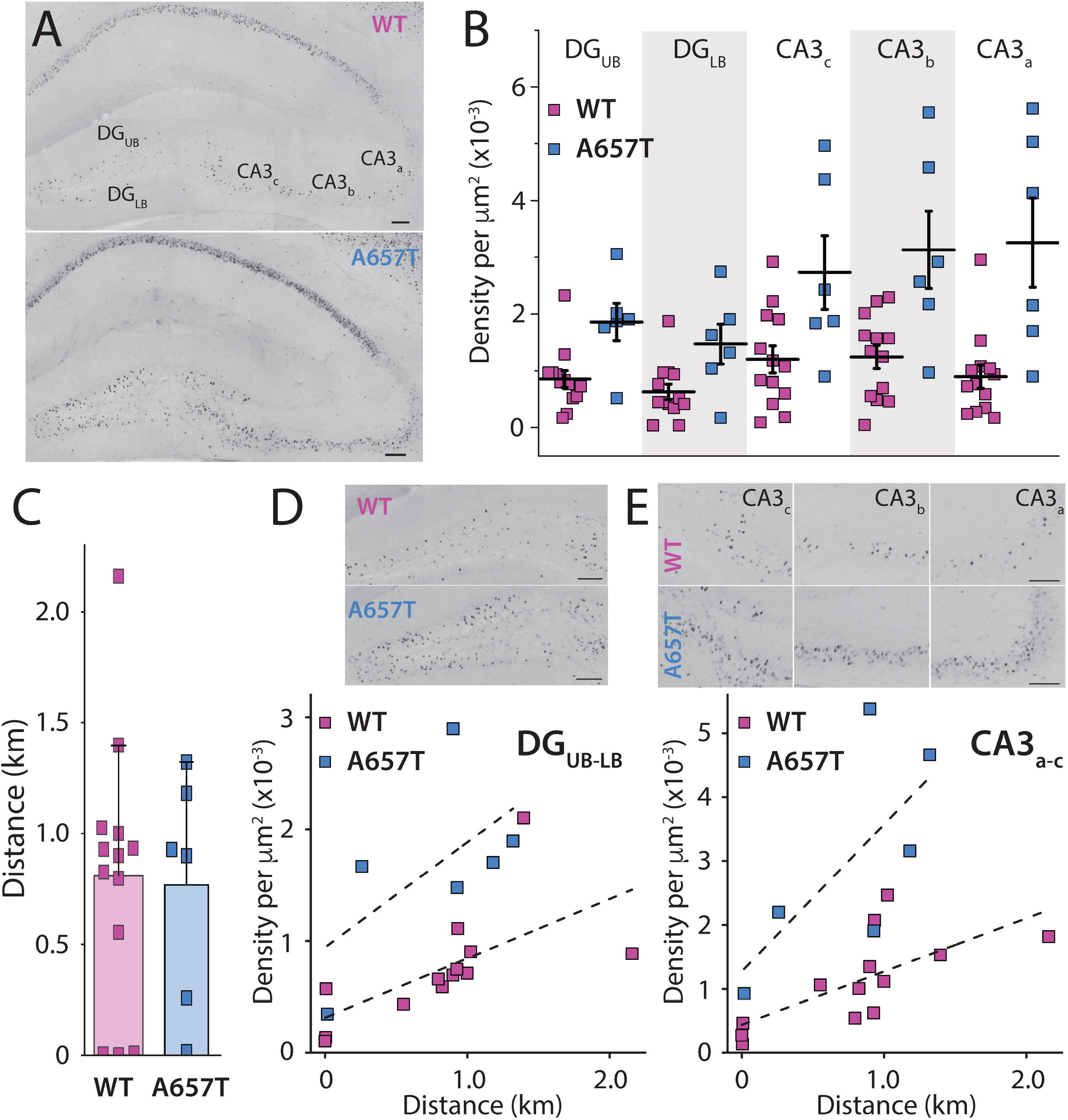
Neuronal activity reporter cFos is elevated in hippocampal neurons in A657T. (**A**) Representative images of the hippocampus after a single bout of running labeled with cFos from WT mice and (**B**) A657T mice. Both example mice had run equivalent distances (WT: 0.898 km; A657T: 0.928 km). cFos cell density was measured in the upper and lower blade of the DG (DG_UB_, DG_LB_) and in each of the subregions of the CA3 a-c. Calibration: 100 µm. (**C**) Total running distance of each animal (**D**) Density of cFos labeled cells in each subregion of the hippocampus and dentate in WT and A657T mice (**E**) Total density of cFos labeled cells of WT and A657T mice as a function of their running distance in the DG and (**F**) CA3 regions.

### EPSP integration and NMDA dependent LTP in distal AC synapses are altered in A657T mice

Our findings demonstrate that neurons in the CA3 region are more active in A657T mice but exhibit only modest changes in their intrinsic properties, suggesting that synaptic mechanisms may play an important role in this hyperexcitability phenotype. We next tested the possibility that CA3 dendritic excitability is elevated in A657T mice, which could enhance the integration and propagation of depolarizing input to distal dendrites. We first determined if spike coupling of AC EPSPs in the *stratum radiatum* was enhanced in A657T mice using minimal stimulation criteria (see methods) of the *stratum radiatum* to evoke four AC EPSPs at 20 Hz. The evoked train of EPSPs, AC1-4, produced a significantly higher spike probability during the train in A657T mice than in WT mice (AC1-4 WT: 26.8 ± 12.6 %, n = 19 cells, 7 mice; AC1-4 A657T: 44.4 ± 20.2 %, n = 13 cells, 6 mice; *p* = 0.039; Mann-Whitney) (Figure 4A). These data are consistent with a generalized increase in dendritic excitability elevating the probability that AC-CA3 synaptic input elicits an AP in CA3 neurons from A657T mice.

**Figure 4:**
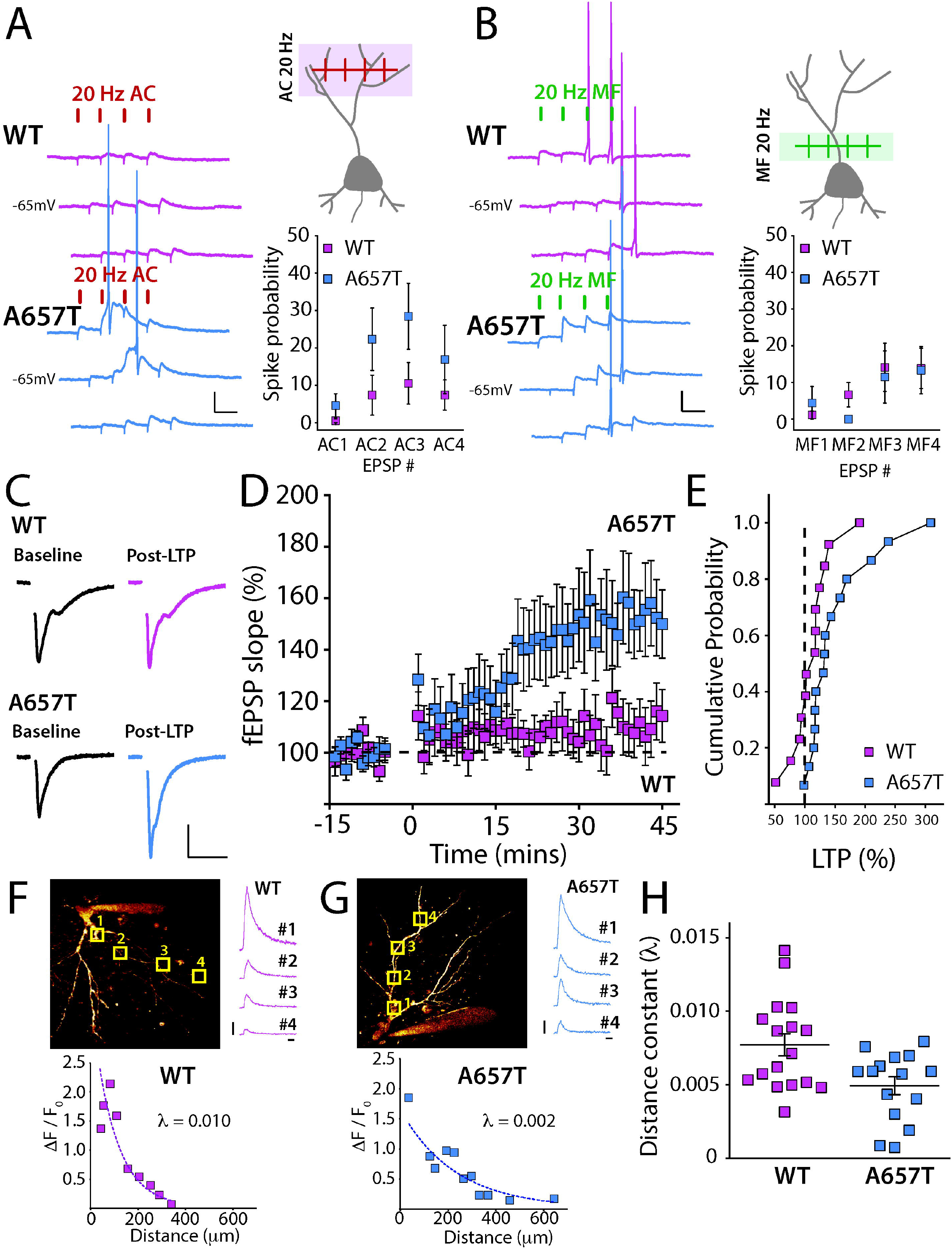
EPSP-spike coupling, bAPs and NMDAR dependent LTP of distal AC synapses is enhanced in A657T mice. (**A**) Representative traces of AC-EPSPs stimulated at 20Hz in CA3 neurons in WT and A657T mice (left). Cartoon representation of AC EPSP stimulation (right, top) and spike probability at each EPSP during the train of stimuli in recordings from WT and A657T mice (right, bottom). Calibration: 10 mV, 50 ms (**B**) Representative MF-EPSPs stimulated at 20 Hz in CA3 neurons (left). Schematic of MF synaptic stimulation (right, top) and spike probability at each MF EPSP during the 20 Hz train in WT and A657T mice (right bottom). Calibration: 10 mV, 50 ms (**C**) Example traces of fEPSPs before and after LTP in WT (top) and A657T mice (bottom). Calibration: 0.2 mV, 20 ms. (**D**) Timecourse of LTP of AC fEPSP responses induced at time = 0 using a 4 x 50Hz burst repeated 10 times at 1 Hz in WT and A657T mice. (**E**) Cumulative distribution for AC LTP in WT and A657T mice. (**F**) 2P-LSM image of the soma and dendrites of a CA3 neuron in slices from WT mice filled with Alexa Fluor 568 through the patch electrode. ROIs are denoted by square boxes as regions of scan sites for action potential evoked Ca^2+^ measurement. Example fluorescence transients measured at ROIs in CA3 neurons in WT mice (right panel). Example recording from a CA3 neuron in WT mice plotting the measured amplitude of Ca^2+^ signal to the distance from soma of the scanning site fit with exponential (bottom panel). (**G**) Image of a CA3 neuron from A657T mice. Fluorescence transients measured at ROIs after somatic depolarization induced APs (right panel). Example recording from experiment from A657T mice (bottom panel). (**H**) Grouped data from all recordings of measured distance constant, λ from exponential fits of dendritic Ca^2+^ evoked by APs in WT and A657T mice.

We next tested whether EPSP-AP coupling of mossy fiber synapses was similarly increased in A657T mice. These synapses are proximal to the soma and therefore changes to dendritic excitability seem less likely to have the same impact on spike coupling as with AC synapses. However, the slowed kinetics of the synaptic KARs at MF synapses (Figure 1J) could affect summation of these inputs in CA3 and produce an analogous change in coupling to AP initiation ^36,37^. We repeated the experiment using minimal stimulation of MF EPSPs to evoke a train of 4 MF EPSPs at 20 Hz or 10 MF EPSPs at 50 Hz (Figure 4B and S2 G & H). In both experiments we found that the AP probability was not different between the two genotypes (MF1-4 20 Hz; WT: n = 18 cells, 11 mice; MF1-4 20 Hz A657T: n = 9 cells, 7 mice; *p* = 0.35 – 1.0 for each bin; Mann-Whitney) (50 Hz; WT: n = 18 cells, 11 mice; A657T: n = 12 cells, 7 mice; *p* = 0.61 – 0.99 for each bin; Mann-Whitney) (Figure 4B and S2 G & H). These observations are consistent with spike coupling of proximal MF synapses not being influenced by dendritic properties, and with prior work suggesting that EPSP-spike coupling in MF is primarily dependent on the large presynaptic dynamics of facilitation of the synapse ^37^, which are not affected in A657T mice (Figure S1 C-F).

The induction of Hebbian synaptic potentiation of distal synapses is affected by active conductances in the dendritic compartment ^38-40^. As there appeared to be a change in the dendritic properties of CA3 neurons in A657T mice, we next tested the prediction that the threshold for NMDA-dependent LTP of AC synapses is reduced by inducing plasticity with a sub-maximal train of stimuli. We recorded AC fEPSPs and induced LTP using a burst of four stimuli at 50Hz with each burst repeated 10 times every second. Consistent with a previous study ^41^, this induction protocol produced a small potentiation in slices from WT mice at 35 - 45 mins after induction (112.1 ± 9.3 %, n = 13 cells, 6 mice) (Figure 4C-E). In contrast, the same stimulation strongly potentiated AC fEPSPs in A657T mice (153.1 ± 15.0 %, n = 15 cells, 7 mice, p = 0.033; Mann-Whitney) (Figure 4C-E). These results suggest that the increased excitability of CA3 neurons, and particularly a potential change in the properties of active dendrites, increases the propensity for Hebbian plasticity of AC synapses in A657T mice.

### CA3 neuron dendritic excitability is increased in A657T mice

Enhanced EPSP-AP coupling of distal AC synapses and a reduced threshold for the induction of NMDA dependent LTP suggests that dendritic excitability is enhanced in CA3 neurons in A657T mice. As an alternate measure of dendritic excitability, we determined the ability of backpropagating action potentials (bAPs) to invade the dendrites of CA3 neurons using two-photon laser scanning microscopy (2PLSM) Ca^2+^ imaging. Neurons were loaded with the Ca^2+^ dye Fluo-4 (200 µM) through the patch electrode and bAPs were evoked by somatic depolarization (Figure 4F & G). The bAP evoked Ca^2+^ signal demonstrated a decremental amplitude at more somatically distal regions of the dendrites (Figure 4 F & G), but in most recordings the Ca^2+^ signal could be detected at least 250 – 300 μm from the soma. A comparison of the decay of the Ca^2+^ signal as a function of the distance from the soma demonstrated that there was less decrement in the signal in recordings from CA3 neurons in the A657T mice (Distance constant λ; WT: 7.71 ± 0.75 x 10^-3^, n = 17 cells, 8 mice; A657T: 4.93 ± 0.61 x 10^-3^, n = 15 cells, 6 mice; *p* = 0.033; Mann-Whitney) (Figure 5 F - H). These results support the conclusion that CA3 neurons in the A657T mice have increased dendritic excitability facilitating the propagation of distal synaptic inputs (and bAPs), which contribute to the enhanced spike transmission and reduced threshold for LTP of AC synapses in CA3 neurons.

**Figure 5:**
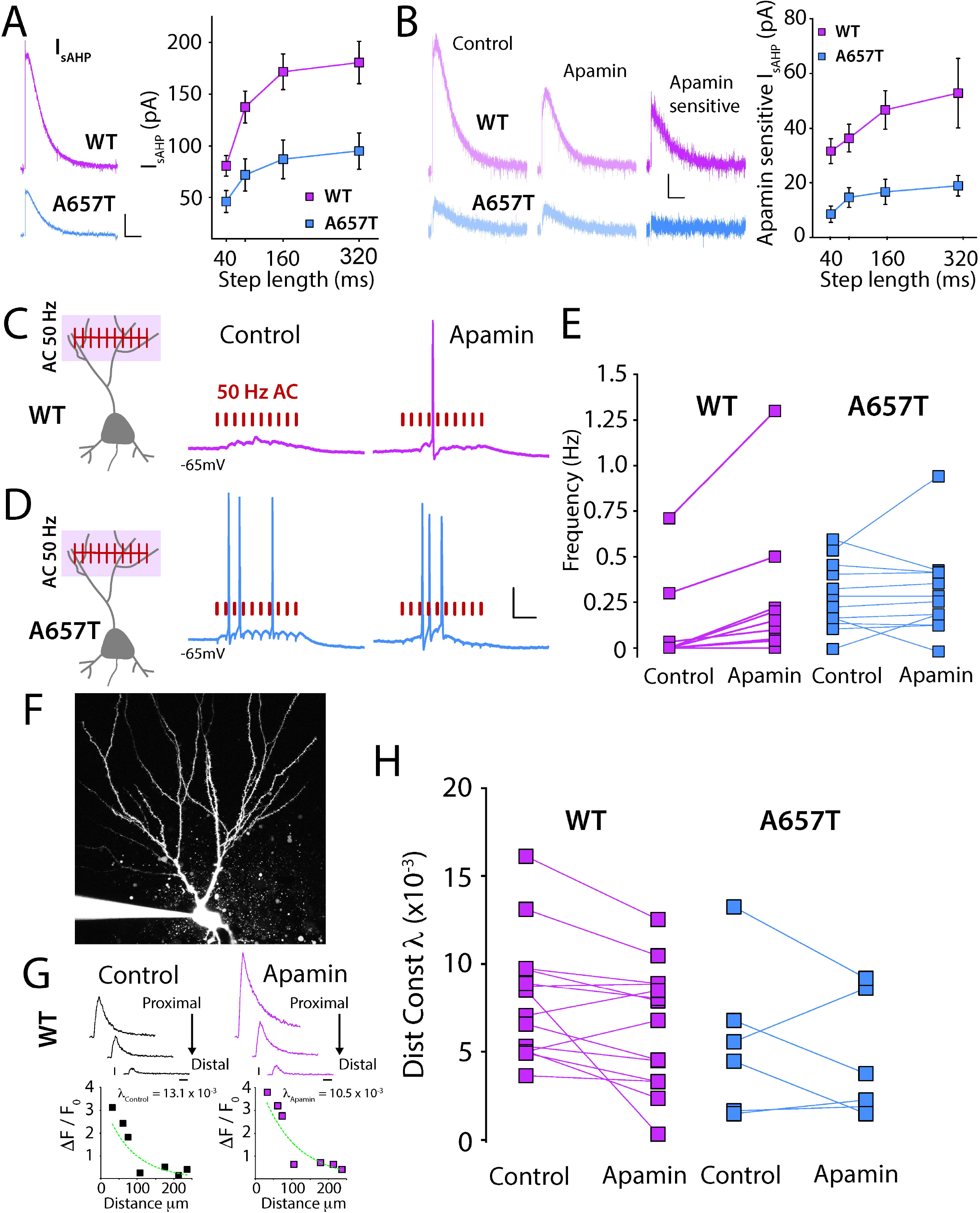
sAHP and SK mediated component of the I_sAHP_ are reduced in A657T mice and effects of apamin on coupling and bAPs. (**A**) Representative currents (I_AHP_) elicited by depolarization in CA3 neurons. Calibration 50 pA, 2.5 s. Amplitude of I*_sAHP_* in CA3 neurons measured with increasing lengths of current injection in WT and A657T mice (right panel). (**B**) Example traces of I*_sAHP_* induced by 160 ms step in CA3 neurons during control, application of apamin (300 nM) and the subtracted trace of the apamin sensitive SK-mediated component from WT mice and A657T mice. Calibration: 25 pA, 2.5 s. Analysis of amplitude of apamin sensitive component of the I*_sAHP_*in CA3 neurons of WT and A657T mice (right panel). I*_sAHP_* was induced using several depolarizing current injections of increasing length. (**C**) Cartoon representation of train stimulation of AC EPSPs and example traces during control and apamin (300 nM) application in WT mice and (**D**) A657T mice. Calibration 20 mV, 50 ms (**E**) Frequency of APs during AC-EPSP train before and after apamin application. (**F**) Example image of the soma and apical dendrites of a CA3 neuron filled with Alexa Fluor 568 through the patch electrode (**G**) Example Ca^2+^ transients measured in the dendrites of WT neuron (top) and all measurements at ROIs at different distances from soma during control and after apamin application. (**H**) Distance constant (λ) calculated from fit of dendritic Ca^2+^ signals measured after evoked APs before and after apamin application in CA3 neurons.

### SK channels underlie the enhanced dendritic excitability and EPSP-spike coupling in A657T mice

A number of K^+^ conductances can limit AP firing and affect dendritic integration of synaptic currents ^42^. As we had found that dendritic excitability was enhanced in A657T mice we focused on K^+^ conductances that mediate the afterhyperpolarization (AHP) in particular, because this conductance can be inhibited acutely by KAR activation ^22,23^ and some of the underlying channels are located in dendrites ^43,44^. The AHP can be measured in the soma following a burst of action potentials and is composed by medium (mAHP) and slow (sAHP) components based on the time course of the potentials. We first measured the somatic AHPs in current-clamp recordings (Figure S3 A-C). There was a significant reduction in the medium component (WT: n = 16 cells, 6 mice; A657T: n = 13 cells, 5 mice; two-way ANOVA, *F*_(1,27)_ = 5.12, *p* = 0.032) (Figure S3B) and the slow component (WT: n = 16 cells, 6 mice; A657T: n = 13 cells, 5 mice; two-way ANOVA, *F*_(1,27)_ = 6.96, *p* = 0.014) of the AHPs in A657T mice (Figure S3C). To confirm that the current underlying the AHP was indeed affected in A657T mice we made voltage clamp recordings and measured the somatic membrane current, I*_sAHP_*which was significantly reduced in A657T mice compared to WT mice (WT: n = 16 cells, 5 mice; A657T: n = 9 cells, 3 mice; two-way ANOVA, *F*_(1,23)_ = 9.50, *p* = 0.0053) (Figure 5A). To test if acute activation of KARs still inhibited the AHP ^22,23^ in mutant mice we tested whether a low concentration of the KAR agonist kainate (50 nM) inhibited both I*_mAHP_* and I*_sAHP_*. The magnitude of the inhibition was not different between genotypes (I*_mAHP_*; WT: 11.7 ± 2.9 %, n = 15 cells, 4 mice; A657T: 32.3 ± 13.2 %, n = 13 cells, 4 mice; *p* = 0.34; Mann-Whitney) (I*_sAHP_*; WT: 26.9 ± 3.9 %, n = 15 cells, 4 mice; A657T: 17.0 ± 9.4 %, n = 13 cells, 4 mice; *p* = 0.088; Mann-Whitney), demonstrating that there is no change in this regulation in the A657T mice.

While there remains some uncertainty about the exact composition of the channels mediating AHPs ^45,46^ prior work has demonstrated that the medium and slow AHP are mediated by Ca^2+^- activated K^+^ channels (SK) in some neurons ^47 48^, including those in the hippocampus ^49^. SK channels are localized to both soma and dendrites of hippocampal pyramidal neurons ^50^, can limit local synaptic depolarizations in CA1 dendrites ^43,51,52^ and constrain NMDA dependent LTP in CA1 Schaffer collateral synapses ^53,54^. We first examined whether the somatically recorded I*_AHP_* in CA3 neurons was sensitive to SK channel blockade. Application of apamin (300 nM), a selective SK channel blocker, significantly inhibited I*_sAHP_*in CA3 neurons in WT mice, demonstrating that a component of the I*_sAHP_*in CA3 is mediated by SK (Figure 5B). Similarly, apamin inhibited the I*_mAHP_*in WT mice (Figure S3 D & E). The sensitivity to apamin of the I*_sAHP_* was significantly diminished consistent with a reduction in the SK contribution to this current in A657T mice (apamin sensitive *I_sAHP_*; WT: n = 19 cells, 7 mice; A657T: n = 13 cells, 5 mice; two-way ANOVA, *F*_(1,30)_ = 10.39, *p* = 0.0030) (Figure 5B). A similar reduction in apamin sensitively was observed in recording of I*_mAHP_* (apamin sensitive *I_mAHP_*; WT: n = 19 cells, 7 mice; A657T: n = 13 cells, 5 mice; two-way ANOVA, *F*_(1,30)_ = 9.97, *p* = 0.0036)(Figure S3 E). These results indicate that SK channels are major constituents of the conductances that underlie the slow and medium components of the somatic I*_AHP_* in CA3 neurons and that activity of this K^+^ channel is significantly diminished in CA3 neurons in A657T mice.

Prior work in CA1 neurons has demonstrated that SK channels in the dendrites are directly responsible for repolarization of glutamate receptor-evoked local dendritic plateau potentials, thereby dampening their propagation ^43^. Therefore, we hypothesized that reduced SK channel activity in CA3 dendrites in A657T mice could be directly responsible for the enhanced propagation of signals from distal synapses. We repeated the AC-spike coupling experiments and tested whether inhibiting the activity of SK channels affected AC-EPSP spike coupling probability. In WT mice, acute application of apamin enhanced the probability of spike transmission after AC synaptic stimulation of 10 minimally evoked EPSPs during a 50 Hz train (WT control: 0.12 ± 0.08 Hz; WT Apamin: 0.28 ± 0.14 Hz; n = 9 cells, 4 mice; *p* = 0.0078; Wilcoxon) (Figure 5 C & E). Thus, inhibiting SK channels mimicked the increased AC-spike coupling phenotype observed in A657T mice. In similar recordings from CA3 neurons in A657T mice we confirmed that AC-spike coupling was enhanced, as we had observed previously, and found that apamin had no detectable effect on AC EPSP spike coupling (A657T control: 0.40 ± 0.01 Hz; apamin: 0.42 ± 0.38 Hz; n = 12 cells, 8 mice; *p* = 0.53; Wilcoxon) (Figure 5D & E), This occlusion of apamin’s effect is consistent with the diminution of SK channel activity in dendrites of CA3 neurons contributing to the observed increase in EPSP-spike coupling of distal AC synapses in A657T mice.

To further assess how SK channel activity affects CA3 dendritic excitability we examined whether apamin affected bAP propagation in CA3 dendrites using 2PLSM Ca^2+^ imaging. In experiments in CA3 neurons from WT animals the decrement of the Ca^2+^ signal along the dendrite was reduced after apamin treatment in the majority of recordings (11 of 14 neurons) (λ _control_: 8.02 ± 0.93 x 10^-3^; λ _apamin_: 6.45 ± 0.92 x 10^-3^; n = 14 cells, 8 mice; *p* = 0.017; Wilcoxon) (Figure 5F - H). This demonstrates that bAP propagation is facilitated in the dendrites of CA3 neurons in the presence of SK channel inhibitor and underlines an important role for SK channels in dampening CA3 dendritic excitability. When we recorded Ca^2+^ signals in CA3 neurons from A657T mice we again found that under control conditions bAPs propagation was enhanced (distance constant λ was reduced) but in this case SK inhibition with apamin had no detectable effect on bAP propagation (λ_control_: 5.60 ± 1.77 x 10^-3^; λ_Apamin_: 4.60 ± 1.42 x 10^-3^, n = 6 cells, 3 mice; *p* = 0.69; Wilcoxon). We also determined whether other conductances that affect dendritic excitability were similarly disrupted in A657T mice. Examination of the K^+^ A-current (I_A_), that is known to affect dendritic excitability of CA1 neurons ^55^ demonstrated that there was no significant change in this conductance in A657T mice (WT: 96.0 ± 1.4 %, n = 16 cells, 5 mice; A657T: 94.8 ± 0.7 %, n = 14 cells, 5 mice, p = 0.82, Mann-Whitney). Taken together these results demonstrate that SK channels that are normally required to dampen dendritic excitability in CA3 neurons are reduced or absent in the mutant A657T mice and thus contribute to enhanced dendritic excitability that affects the integration of synaptic signals and the induction of LTP in AC synapses.

## Discussion

### A657T mutation in GluK2 receptor affects dendritic excitability through a downregulation of SK channels

We have discovered a surprising cellular and synaptic pathophysiology caused by the introduction of a disease-causing missense mutation in the *Grik2* gene. The most prominent finding is that dendritic excitability is increased through downregulated SK channel function in KAR mutant CA3 neurons. The modulatory activity of KARs on AHPs in hippocampal neurons has been explored over many years ^21-23^. For example, *in vitro* recordings in CA3 neurons demonstrated that induction of long-term depression of KARs relieves activity-dependent inhibition of I*_sAHP_* thus reducing neuronal excitability and supporting the idea that I*_sAHP_* is tonically modulated by KARs ^56^. KARs therefore provide a feedback mechanism linking the activity of iGluRs to neuronal excitability through the K^+^ conductances that underlie AHPs. These same channels in many instances play a significant role in regulating dendritic excitability. The relevance of this interplay between synaptic and intrinsic conductances to hippocampal circuits *in vivo* remains unclear because prior evidence for this modulatory activity was limited to *in vitro* studies. Here, while assessing the effect of a missense mutation that affects gating of GluK2 containing receptors in a newly developed mouse model, we unexpectedly found that CA3 neurons were much more active in slices from A657T mice.

While this activity could in principle be due to an elevated intrinsic propensity for spiking, our findings suggest that the elevations in AP frequencies are synaptically driven. Intrinsic excitability was not grossly altered, and the I-O curve of somatically evoked APs was normal. In contrast there was a clear elevation in the probability of EPSP-spike coupling in distal synapses. A likely scenario, therefore, is that the observed elevation in neuronal activity was because of an elevation in the integration of synaptic inputs due to an increase in dendritic excitability in A657T mice. This was supported by experiments blocking SK channels, known to regulate dendritic excitability, which replicated the phenotype in WT neurons but had little effect in A657T animals. Thus, our findings demonstrate that SK channel function is decreased, which enhances dendritic integration and results in more active CA3 neurons in A657T mice.

The excitability of dendrites of hippocampal and cortical neurons are regulated by the localization of several voltage gated and voltage independent ion channels that impact the propagation of synaptic depolarizations ^43,55,57,58^. Small conductance Ca^2+^ activated channels have been immunohistochemically localized to dendrites of many neuron types, including CA3 pyramidal neurons in the hippocampus ^50,59^. Moreover, in CA1 neurons SK channels are active in distal dendrites and limit the spread of synaptic depolarizations ^43^. However, decreased SK mediated conductance and an associated increase in excitability of dendrites that elevated EPSP-AP coupling in A657T mice was unexpected. This potentially provides an unexpected cellular phenotype that could have a profound effect on circuit function beyond a kinetic change in synaptic KARs that makes a relatively modest postsynaptic contribution in CA3 synapses. For instance, SK channel inhibition increases the power of gamma oscillations in the CA3 ^60^ which could have significant effects on cognition ^61^.

SK channels can also limit synaptic plasticity induction. Apamin blockade of SK channels reduces the threshold for NMDA dependent LTP in CA1 neurons ^54^. We found that a relatively short induction train ^41^ caused minimal potentiation (∼ 10 %) in slices from WT animals but an exaggerated potentiation (∼ 50 %) in A657T mice. These results suggest that, as in CA1 neurons ^53,54,62^, SK channels localized to dendrites and spines act as a brake to limit LTP of distal CA3 AC synapses, the regulation of which is lost in A657T mice. Hippocampal LTP is intimately linked with memory formation and any perturbation of memory processes is likely to have behavioral consequences by disrupting cognition. While we do not report on behavior in this study, we did measure expression of the immediate early gene cFos as an *in vivo* proxy for the activity of neurons, providing a snapshot of the relative amount of activity. We found that cFos expression was elevated in all hippocampal subregions in A657T mice providing strong support for the conclusion that neurons are more easily activated *in vivo* in these animals.

### Dendritic excitability and neurodevelopmental disorders

An emerging theme in establishing the cellular correlates of circuit dysfunction in models of neurodevelopmental disorders has focused on synaptic mechanisms including receptors, scaffolds and cell adhesion molecules that have been directly implicated in disease ^63 64,65^. These synaptic deficits in many cases are primary causes of changes in the balance of excitatory and inhibitory synaptic transmission which are associated with many neurodevelopmental and neuropsychiatric disorders ^66^. Similarly, there are numerous channelopathies that are caused by variants in both voltage gated and Ca^2+^ gated ion channels ^67,68^. Despite this there are only a few examples of studies that have directly assessed dendritic mechanisms in mouse models of NDDs and yet fewer studies in which dendritic excitability has been characterized in detail. As one example, ASD can result from loss-of-function mutations in the *SCN2A* gene, which encoded the Nav1.2 sodium channel ^69^. *Scn2A* haploinsufficiency in mice causes a reduction in excitability of the dendritic compartment of cortical pyramidal cell dendrites, reducing bAPs and synaptic plasticity ^70^. In the *Fmr1* KO mouse, the model of fragile X syndrome, multiple ion channels have been reported to be disrupted and in CA1 neurons a specific loss of dendritic Kv4 elevated the excitability of the dendritic compartment, increased propagation of bAPs and enhanced synaptic plasticity ^71^. These studies demonstrate that either direct or indirect loss of voltage dependent channels in the dendritic compartment can affect dendritic computations in mouse models of NDD.

In our study a missense mutation in the gene encoding the glutamate gated receptor subunit GluK2 revealed a functional coupling *in vivo* to SK channels that regulate dendritic excitability. The relevance of KAR modulation of AHPs to brain function has been a mystery, therefore this study provides evidence that this functional coupling could play a significant role in the normal physiological properties of CA3 neurons. Aberrant SK activity has also been described in other NDD mouse models. Haploinsufficient *Pten* mice, another ASD model, have a marked increase in SK function in L2/3 cortical neurons causing a decrease in excitability but not affecting dendritic integration of subthreshold synaptic events ^72^. In *Fmr1* KO mice, it was recently demonstrated that SK currents are reduced in CA3 neurons because of a loss of a direct protein interaction between Fmrp and SK2 causing elevated neuronal excitability ^73^. It remains to be tested whether CA3 dendritic excitability and synaptic integration are affected in *Fmr1* KO mice similar to what we have observed in the A657T mice. Conversely, loss of *Ube3a* ubiquitination of SK2 in the mouse model of Angelman syndrome results in increased spine expression of SK2 in CA1 neurons and impairs LTP induction ^74^. Thus, there is a growing appreciation that SK dysfunction in mouse models of NDDs can disrupt excitability and synaptic plasticity mechanisms. The importance of SK to normal brain development and function is further highlighted by the recent discovery of *de novo* and inherited loss of function variants in *KCNN2*, which encodes the SK2 channel, in patients with delays in motor, intellectual and language development ^75^. These phenotypes in SK2 disorders overlap with the clinical phenotype observed in the individuals carrying the A657T mutation in *GRIK2* gene ^6,7^.

### Limitations of the Study

The mechanistic basis for alterations in K^+^ channel activity and dendritic excitability caused by this particular *Grik2* mutation and disruption of the channel pore region is not clear. Unfortunately, our proteomic analysis did not detect Kcnn2 peptides (SK2 channel) that could be quantified, which is often the case for low expression proteins, and we may require other methodologies for future assessment of SK channel expression and localization. These effects on SK are likely pleiotropic due to developmental consequences that cannot be extricated from their acute effects on the phenotype. Indeed, in this case we are modeling a neurodevelopmental disorder in humans and would expect the mutation to have effects throughout the life of the animal. We did in fact see elevated spiking of CA3 neurons in perinatal mice (P12 – P15; data not shown), similar to the adult phenotype we report in the paper, suggesting that this is also prevalent when the brain is still developing. Future mechanistic studies using genetic or viral tools to conditionally express mutant receptors could shed further light on these questions.

There is a growing list of variants in glutamate receptor genes associated with human NDDs, but it is not at all clear how those mutations cause functional alterations of neural circuits. Here we found that an orthologous single amino acid mutation in the GluK2 in a mouse had an unexpected effect on K^+^ channel function and dendritic excitability. Our findings provide novel insight into how disease-causing iGluR mutations can disrupt neuronal and circuit function.

## Methods

### Animal care and use

All experimental procedures were conducted in accordance with the ethical policies and protocols approved by the Northwestern University IACUC. The point mutation was engineered into the genome of the mouse using the CRISPR/Cas9 gene editing system to generate the A657T amino acid substitution in the mouse *Grik2* gene. Single guide RNAs (gRNA), Cas9 mRNA and a single strand oligonucleotide donor (ss donor) with a single base mutation were microinjected into mouse zygotes. Founders with the positively identified A657T mutation were bred with wild type C57/bl6J mice, and each of these founders produced viable heterozygous mutant mice. Successful generation of the founders and offspring were confirmed by direct sequencing. Heterozygous mutant male mice were crossed with WT female mice for breeding, and both male and female WT and heterozygous A657T littermates were used for experiments. Mice were housed with food and water provided *ad libitum* under 12/12 hours dark-light cycle. Experiments were conducted with the investigator blind to the genotype of the animals. These were subsequently confirmed by tail biopsy samples via post hoc sequencing and/or PCR.

### Electrophysiology

Male and female GluK2 WT or A657T heterozygous mice (3 – 5 weeks old) were deeply anesthetized with inhalation of isoflurane and an intraperitoneal injection of xylazine (10 mg/kg) and ketamine (100 mg/kg). Mice underwent trans-cardiac perfusion with an ice-cold sucrose artificial cerebrospinal fluid (ACSF) solution containing (in mM): 85 NaCl, 2.5 KCl, 1.25 NaH_2_PO_4_, 25 NaHCO_3_, 25 glucose, 75 sucrose, 0.5 CaCl_2_, and 4 MgCl_2_, including 10 μM DL-APV and 100 μM kynurenate. Horizontal brain sections (350 μm-thick) were prepared in the same ice-cold sucrose ACSF on a Leica Vibratome (Leica Microsystems, Inc). Slices were incubated in the same sucrose ACSF at 30 – 32°C for ∼30 min, then the solution was gradually exchanged for a recovery ACSF containing (in mM): 125 NaCl, 2.4 KCl, 1.2 NaH_2_PO_4_, 25 NaHCO_3_, 25 glucose, 1 CaCl_2_, and 2 MgCl_2_, including 10 μM DL-APV and 100 μM kynurenate at room temperature. Slices were transferred to a recording chamber after a recovery period of at least 1.5 h and were visualized under Dodt-Gradient-Contrast optics (Luigs & Neumann). During recordings, slices were perfused with normal ACSF (28 – 30 °C) containing (in mM): 125 NaCl, 2.4 KCl, 1.2 NaH_2_PO_4_, 25 NaHCO_3_, 25 glucose, 2 CaCl_2_, and 1 MgCl_2_ continuously equilibrated with 95 % O_2_ and 5 % CO_2_. Standard techniques were used to make patch-clamp recordings from visually identified CA3 pyramidal neurons in the hippocampus. Recording electrodes had tip resistances of 2 – 4 MΩ when filled with a cesium-based internal solution containing (in mM): 95 CsF, 25 CsCl, 10 Cs-HEPES, 10 Cs-EGTA, 2 NaCl, 2 Mg-ATP, 10 QX-314, 5 TEA-Cl, and 5 4-AP for voltage clamp recordings or a potassium-based internal solution containing (in mM): 125 KMeSO_4_, 5 KCl, 5 NaCl, 0.02 EGTA, 11 HEPES, 1 MgCl_2_, 10 phosphocreatine, 4 Mg-ATP, 0.3 Na-GTP for current clamp recordings unless further specified. Liquid junction potential was not corrected. In voltage clamp access resistance (R_a_) was continuously monitored, and recordings were discarded when R_a_ showed >20% change during the experiments. For voltage clamp recordings cells were held at -70 mV unless otherwise specified. Data were acquired and analyzed using a Multiclamp 700B amplifier and pClamp 10 software (Molecular Devices).

Synaptic afferents were stimulated with a monopolar glass electrode filled with normal ACSF positioned in the stratum lucidum or in the stratum radiatum to elicit mossy fiber (MF) or associational-commissural (AC) mediated synaptic responses. EPSCs were isolated using the GABA_A_ receptor antagonists bicuculline (10 μM) and picrotoxin (PTX) (50 μM). MF-EPSCs were characterized by large short-term synaptic plasticity, rapid rise-time kinetics, and when appropriate the Group II mGluR agonist DCG-IV was applied at the end of recordings ^76 77 78 79^. Kainate receptor mediated EPSCs (EPSC_KAR_) were isolated using the AMPA receptor antagonist GYKI53655 (50 μM). EPSC decay kinetics were quantified by fitting the traces with single or double exponential. For MF-LTP recordings, repeated tetanic stimuli composed of 100 bursts at 100 Hz repeated three times every 20 second were applied after obtaining a 10 min stable baseline. IPSCs were recorded in the presence of AMPA and kainate receptor blockers CNQX (10 μM) and an NMDA receptor blocker D-APV (50 μM).

For AC-LTP experiments field EPSPs (fEPSPs) were measured using extracellular recordings. Recording electrodes had tip resistances of 2 – 4 MΩ when filled with normal ACSF. Stimulating and recording electrodes were placed in the stratum radiatum. The slope of the fEPSPs was measured for LTP recordings. After obtaining a stable baseline for at least 10 min, PTX was perfused into the recording solution for 5 min to block GABA_A_ receptors. Short burst high frequency stimuli (four bursts at 50Hz repeated 10 times every second) was applied to induce LTP of AC-fEPSPs ^41^.

Spontaneous action potential firing was measured in cell attached recordings of action currents in voltage clamp mode. For cell attached experiments, the recording electrodes had tip resistances of 3 – 5 MΩ when filled with the same potassium-based internal solution as for current clamp recordings. The effect of NBQX on firing frequency was analyzed by a paired comparison in each cell between baseline and NBQX-treated conditions.

Intrinsic properties were analyzed in current clamp recordings as previously described ^80^. Resting membrane potential was determined by switching the recording mode to current clamp immediately after forming the whole cell configuration. Rheobase was measured by gradually increasing the size of injected depolarizing square currents in 5 pA increments until the first AP was elicited. AP threshold was considered to be the potential where the first derivative of the AP (*dV/dt*) showed an abrupt upward inflection when *dV/dt* was plotted against the corresponding membrane voltage. Input-output frequency response curves for APs were generated by injecting depolarizing square currents (0 – 1000 pA with 100 pA increments, 500 ms width) to analyze AP firing properties. Baseline membrane voltage was adjusted to -65 mV by injecting continuous currents through the patch pipettes.

Voltage-dependent conductances were analyzed in voltage clamp recordings using a K^+^ based internal solution. For recordings of total Na^+^ (I_Na+total_) and K^+^ (I_K+total_) conductances, membrane potential was clamped, and depolarizing step currents (500 ms) were injected from -70 mV to +30 mV (5 mV increments). I_Na+total_ could be detected starting at ∼ -50 - -40 mV in most cells. I_K+total_ was measured by averaging the plateau current of the last 5 ms during the voltage steps. Voltage dependent fast K^+^ current (A-current; I_A_) was measured in normal ACSF but concentrations of Ca^2+^ and Mg^2+^ were modified to 0 and 3 mM, respectively, to block Ca^2+^ mediated and activated currents. The Na^+^ channel blocker tetrodotoxin TTX (0.5 μM) and the K^+^ channel blocker tetraethylammonium (TEA) (1 mM) were included to the recording solution to block Na^+^ currents and TEA-sensitive K^+^ currents. Cells were clamped at -80 mV and depolarized to +20 mV with a depolarizing voltage step (200 ms) followed by an identical voltage step 100 ms after the 1^st^ step to activate voltage dependent K^+^ channels. I_A_ was measured as the percentage of the peak current evoked by the 2^nd^ voltage step relative to the current evoked by the 1^st^ step, reflecting fast recovery of the channels from inactivation ^81^.

EPSP-AP coupling was analyzed using stimulation of EPSP that was termed minimal using the following experimental criteria. After obtaining a recording of an EPSP stimulation intensity was reduced to the minimum current amplitude at which it was possible to evoke a synaptic response with no observed failures. This resulted in starting EPSPs that were not different between genotypes for either MF or AC EPSPs. Baseline membrane voltage was adjusted to - 65 mV by injecting continuous currents through the recording electrodes. Spike probability in each stimulation bin was measured in ∼10 sweeps and was then averaged in each cell.

AHPs and I*_AHP_* were induced by depolarizing currents injection in current clamp and voltage clamp recordings, respectively. In current clamp recordings, the baseline voltage was maintained at -65 mV with continuous current injection. Depolarizing square current injection (0 – 1000 pA with 100 pA increments, 500 ms width) was used to induce AHPs. mAHP and sAHP were measured at their peaks typically several tens of ms (mAHP) and several hundreds of ms (sAHP) ms after the depolarizing step. When the peaks were not clearly distinguishable, mAHP and sAHP were measured at 50 – 100 ms (mAHP) and 500 – 1000 ms (sAHP) after the depolarization. In voltage clamp recordings, I*_mAHP_* and I*_sAHP_*were evoked by depolarizing the cells from -60 mV to +60 mV with square pulses (40, 80, 160, and 320 ms) and peaks measured typically at several tens of ms for I*_mAHP_* and several hundreds of ms for I*_sAHP_* after the depolarization. When the peak currents were not clearly delineated the, I*_mAHP_* and I*_sAHP_* were measured 20 – 100 ms (I*_mAHP_*) and 150 – 1150 ms (I*_sAHP_*) after the depolarization.

### Ca^2+^ imaging

Two-photon laser scanning microscopy (2PLSM) images of CA3 pyramidal neurons in the hippocampus were obtained using methods like those previously described ^80,82,83^. Cells were loaded with Alexa Fluor 568 (50 μM) through the patch pipette for visualization. The dye was perfused intracellularly for at least 20 min after forming the whole cell configuration. Images were acquired using a femtosecond pulsed excitation at 790 nm (Mira 900P with a Verdi 10W pump, Coherent Laser). Laser power was controlled with an M350 (KD*P) series Pockels cells (ConOptics). A Prairie Ultima (Bruker Nano, Fluorescence Microscopy Unit) scan head on an Olympus BX-61 upright microscope was used for imaging the slice with two Hamamatsu R3982 side on photomultiplier tubes. For Ca^2+^ imaging, cells were filled with the Ca^2+^ indicator Fluo-4 (200 μM) loaded through patch pipettes. Dendritic Ca^2+^ signals were triggered by 10 backpropagating APs (bAPs) evoked by somatic suprathreshold depolarizing current injection (1 nA for 1 ms at 100 Hz). Ca^2+^ signals in multiple (typically 8 - 10) regions of interests (ROIs) within a single branch of dendrites were acquired as fluorescent line scan signals in 3 – 5 pixels (0.77 μm) per line with 22.8 μs dwell time. The decay of the Ca^2+^ signals along the dendrite were fit using an exponential equation (*y = Ae^-λx^)* to derive the distance constant (λ) as a proxy of dendritic excitability. All components were configured and acquired using Multiclamp 700B amplifier and Prairie View 9.0 software (Bruker).

### Wheel Running

A wireless low profile running wheel (Med Associates) was placed into the homecage of single housed mice during the dark cycle. Mice were allowed free access to the wheel for 90-120 minutes and running activity logged for each animal. Mice were sacrificed and brain removed for cFos immunohistochemistry.

### Immunohistochemistry

Immunohistochemistry for GluK2/3 was carried out according to prior published methods ^84^. Briefly, mice were anesthetized with isoflurane, decapitated, brains removed and rapidly frozen in liquid nitrogen or dry ice. Brains were sectioned coronally on a cryostat (50μm), fixed in modified Carnoy’s solution (1:6, acetic acid:ethanol), washed in PBS, permeabilized in 0.1% TritonX100 and 0.05% Tween20, blocked in 5 % newborn calf serum with 0.05% Tween20, and incubated overnight at 4°C in Rabbit anti-GluK2/3, (1:100; Invitrogen PA1-37780) and Mouse anti-NeuN, (1:250; Millipore MAB377). The following day the sections were washed in PBS and incubated with Goat anti-Rabbit-488 and Goat anti-Mouse-568 (1:200) for 2 hours, washed with PBS and coverslipped with ProLong Diamond mounting media containing Dapi (ThermoFisher Scientific).

Immunohistochemistry for cFos was carried out according to the general method described previously ^31^. Briefly, mice were anesthetized with isoflurane and perfused through the heart with PBS containing 0.02 % sodium nitrite, followed by 2% paraformaldehyde (PFA) in 0.1M sodium acetate buffer (pH 6.5) and 2% PFA in 0.1 M sodium borate buffer (pH 8.5). The brains were removed from the skull, post-fixed for 3 hours and sectioned on a vibrating microtome (Leica Vibratome, VT1000). Free-floating, coronal sections (50μm) were washed in TBS containing 0.01% H_2_O_2_, permeabilized in TBS containing 0.001% Triton X-100 and 0.001% NP40 and blocked for 1 hour in TBS containing 0.05% bovine serum albumin (TBS-BSA), 1% normal donkey serum (NDS) and unconjugated Donkey anti-Rabbit IgG (1:1,000). Sections were then washed in TBS-BSA and incubated overnight on a shaker in TBS-BSA containing Rabbit anti-cFos (1:2,000; Cell Signaling Product# 2250).

The following day the sections were washed in TBS-BSA and incubated in TBS-BSA, 0.25% NDS and Donkey anti-Rabbit IgG-biotin (1:1,000) for 1 hour. The sections were then washed in TBS and incubated for 1 hour in TBS containing streptavidin-biotin-peroxidase complex (ABC 1:1,000 TBS; ThermoFisher Scientific, Product# 32020), washed again in TBS, and incubated in TBS containing diaminobenzidine. Sections were mounted onto slides, air-dried overnight, dehydrated in ethanol, xylene, and coverslipped using Permount.

### Synaptosome preparation and Protein Gels

P3 synaptosomes containing the large mossy fiber fraction were prepared as previously described ^31^. Briefly, hippocampi isolated from adult mice were homogenized in 0.325 M sucrose and then centrifuged at 900 xG for 10 min at 4°C. The supernatant containing the small synaptosomes (P2) was collected, and the remainder resuspended in 0.325 M sucrose and centrifuged at 900 xG for an additional 10 min at 4°C. The pooled supernatants were then centrifuged at 17,000 × G for 55 min at 4°C. The resultant pellet is enriched with small hippocampal synaptosomes (P2). The large mossy fiber synaptosomes sedimented with the nuclei after the initial low-speed spin and were resuspended in 1.5 ml of 18% Ficoll in 0.325 M sucrose and centrifuged at 7500 × G for 40 min. The supernatant from this spin was collected, diluted in 4 volumes 0.3 M sucrose, and centrifuged at 13,000 × G for 20 min at 4°C to yield the P3 pellet containing the large mossy fiber synaptosomes.

Protein isolated from the small hippocampal synaptosomes (P2) and the large mossy fiber synaptosomes (P3) was diluted in 0.325 M sucrose and loading buffer (50 mm Tris-Cl, 100 mm β-mercaptoethanol, 2% SDS, 0.1% bromophenol blue, and 10% glycerol) and run on a 4-12% gradient polyacrylamide gel. Gels were transferred to PVDF membrane, blocked in 20mM TBS containing 3% IgG-free BSA and 1% normal donkey serum, and blotted overnight in the same solution containing the following antibodies: mouse anti-β-actin (1:30,000), Rabbit anti-GluK2/3 (1:1,000), and mouse anti-postsynaptic density 95 (PSD95) (1:5,000). The following day blots were washed 3x 5 minutes in TBS and incubated for 1 hour in the blocking solution (above) containing both secondary antibodies (Donkey anti-Rabbit IgG, Donkey anti-Mouse IgG) and visualized on a Li-Cor Odyssey Fc fluorescent imager.

### Sample preparation for Mass spectrometry

Tissue from the CA3 region of the hippocampus was micro-dissected from WT and A657T Het mice and homogenized in RIPA lysis buffer (50 mM Tris base, 150 mM NaCl, 1% Triton X-100, 2 mM EDTA, 1% sodium deoxycholate, and 1% SDS with complete Mini, EDTA-free protease inhibitor cocktail) using the Kimble Pellet Pestle cordless motor and grinders. Homogenates were rotated at 4°C for 30 min and pelleted by centrifugation for 15 min at 14,000 *g*. The supernatant was separated into a new tube, precipitated using the chloroform/methanol method, denatured with 8 M urea (in a 100 mM ammonium bicarbonate [ABC] vortex 1 h at RT) and processed with ProteaseMAX according to the manufacturer’s protocol. The samples were reduced with 5 mM Tris(2-carboxyethyl)phosphine (TCEP; vortexed for 1 h at RT), alkylated in the dark with 10 mM iodoacetamide (IAA; 20 min at RT), diluted with 100 mM ABC, and quenched with 25 mM TCEP. Samples were sequentially digested with Lys-C (2 h at 37°C with shaking) and trypsin (overnight at 37°C with shaking), acidified with trifluoroacetic acid (TFA) to a final concentration of 0.1%, and spun down (15,000 *g* for 15 min at RT) wherein the supernatant was moved to a new tube. The samples were desalted using Peptide Desalting Spin Columns (89852; Pierce) and dried down with vacuum centrifugation. The samples were resuspended in 100 mM HEPES, pH 8.5, Micro BCA (23235, Pierce) assay was performed to determine the peptide concentration, and 95 μg of each sample was used for isobaric labeling. TMT labeling was performed on the samples according to the manufacturer’s instructions (A44520; Thermo Scientific). After incubation for 2 h at RT, the reaction was quenched with 0.3% (v/v) hydroxylamine, isobaric-labeled samples were combined 1:1:1:1:1:1:1:1:1:1, and the pooled sample was dried down with vacuum centrifugation. The sample was resuspended in 0.1% TFA solution, desalted using Peptide Desalting Spin Columns (89852; Pierce), and dried down with vacuum centrifugation. The sample was resuspended in 0.1% TFA solution and fractionated into eight fractions using a High pH Reversed-Phase Peptide Fractionation Kit (84868; Pierce) wherein fractions were step eluted in 300 μl of buffer of increasing acetonitrile concentrations with decreasing concentrations of 0.1% triethylamine per the manufacturer’s instructions (5.0%, 10.0%, 12.5%, 15.0%, 17.5%, 20.0%, 22.5%, 25.0% and 50% of ACN in

0.1% triethylamine solution). The fractions were dried down with vacuum centrifugation. The high pH peptide fractions were directly loaded into the autosampler for MS analysis without further desalting.

### Mass spectrometry

Two micrograms of each fraction or sample were auto-sampler loaded with an UltiMate 3000 HPLC pump onto a vented Acclaim Pepmap 100, 75 µm x 2 cm, nanoViper trap column coupled to a nanoViper analytical column (Thermo Fisher Scientific, Cat#: 164570, 3 µm, 100 Å, C18, 0.075 mm, 500 mm) with stainless steel emitter tip assembled on the Nanospray Flex Ion Source with a spray voltage of 2000 V. An Orbitrap Fusion (Thermo Fisher Scientific) was used to acquire all the MS spectral data. Buffer A contained 94.785% H_2_O with 5% ACN and 0.125% FA, and buffer B contained 99.875% ACN with 0.125% FA. For TMT MS experiments, the chromatographic run was for 4 hours in total with the following profile: 0-7% for 7, 10% for 6, 25% for 160, 33% for 40, 50% for 7, 95% for 5 and again 95% for 15 mins receptively.

We used a multiNotch MS3-based TMT method to analyze all the TMT samples ^85,86^. The scan sequence began with an MS1 spectrum (Orbitrap analysis, resolution 120,000, 400-1400 Th, AGC target 2×10^5^, maximum injection time 200 ms). MS2 analysis, ‘Top speed’ (2 s), Collision-induced dissociation (CID, quadrupole ion trap analysis, AGC 4×10^3^, NCE 35, maximum injection time 150 ms). MS3 analysis, top ten precursors, fragmented by HCD prior to Orbitrap analysis (NCE 55, max AGC 5×10^4^, maximum injection time 250 ms, isolation specificity 0.5 Th, resolution 60,000).

### MS data analysis and quantification

Protein identification/quantification and analysis were performed with Integrated Proteomics Pipeline - IP2 (Bruker, Madison, WI. http://www.integratedproteomics.com/) using ProLuCID ^87,88^, DTASelect2 ^89,90^, Census and Quantitative Analysis (For TMT MS experiments). Spectrum raw files were extracted into MS1, MS2 and MS3 (For TMT experiments) files using RawConverter (http://fields.scripps.edu/downloads.php). The tandem mass spectra were searched against UniProt mouse protein database (downloaded on 10-26-2020) ^91^ and matched to sequences using the ProLuCID/SEQUEST algorithm (ProLuCID version 3.1) with 5 ppm peptide mass tolerance for precursor ions and 600 ppm for fragment ions. The search space included all fully and half-tryptic peptide candidates within the mass tolerance window with no-miscleavage constraint, assembled, and filtered with DTASelect2 through IP2. To estimate peptide probabilities and false-discovery rates (FDR) accurately, we used a target/decoy database containing the reversed sequences of all the proteins appended to the target database ^92^. Each protein identified was required to have a minimum of one peptide of minimal length of six amino acid residues; however, this peptide had to be an excellent match with an FDR < 1% and at least one excellent peptide match. After the peptide/spectrum matches were filtered, we estimated that the peptide FDRs were ≤ 1% for each sample analysis. Resulting protein lists include subset proteins to allow for consideration of all possible protein forms implicated by at least three given peptides identified from the complex protein mixtures. Then, we used Census and Quantitative Analysis in IP2 for protein quantification of the TMT MS experiment. TMT MS data were normalized using the built-in method in IP2.

Spyder (MIT, Python 3.7, libraries, ‘numpy’, ‘scipy’) was used for data analyses. RStudio (version, 1.2.1335, packages, ‘tidyverse’) was used for data virtualization. The Database for Annotation, Visualization and Integrated Discovery (DAVID) (https://david.ncifcrf.gov/) was used for protein functional annotation analysis.

#### QUANTIFICATION AND STATISTICAL ANALYSIS

Statistical analyses were done using GraphPad Prism (GraphPad Software), and Origin (OriginLab) software. Comparisons were made using Mann-Whitney U test or the Kolmogorov-Smirnov (K-S) test for unpaired two samples. Paired two samples were compared using Wilcoxon signed-rank test. For multiple comparisons repeated two-way ANOVA followed by *post hoc* Bonferroni’s correction was employed. Differences were considered to be significant when *p* < 0.05. Data are presented as mean ± SEM.

## Acknowledgements

This work was supported by NIH/NINDS R01NS105502 to GTS and AC and NIH/NINDS R01NS115471 to AC. Mice were generated with the assistance of Northwestern University Transgenic and Targeted Mutagenesis Laboratory.

## Figure Legends

**Figure S1:**
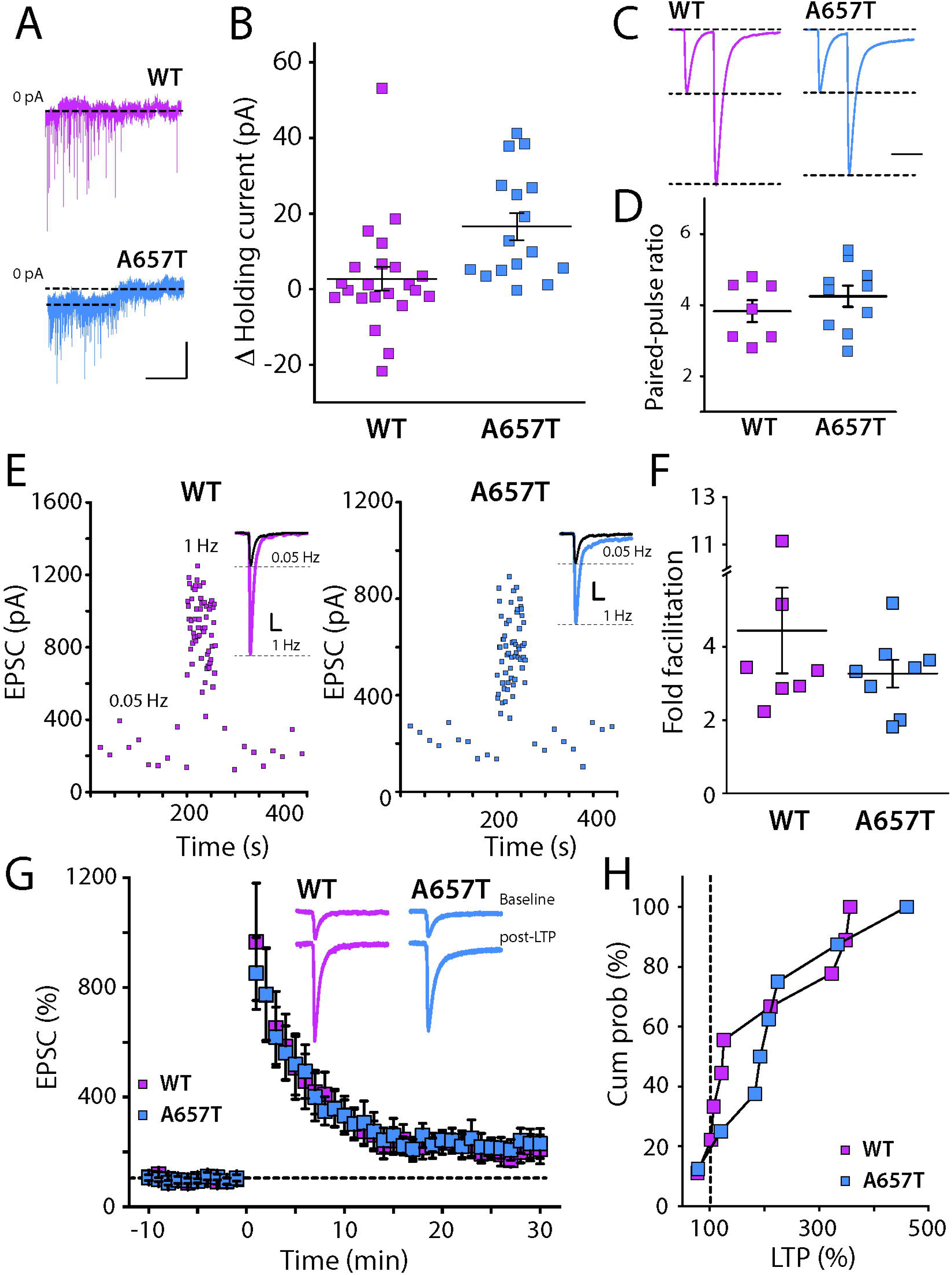
Tonic KAR mediated current and presynaptic mossy fiber synaptic plasticity in A657T mice. **(A**) Example traces of voltage clamp recordings from CA3 neurons during application of NBQX (50 µM). Calibration: 50 pA, 60 s (**B**) Change in holding current of CA3 neurons after NBQX in WT and A657T mice (**C**) Example traces of pairs of mossy fiber EPSCs stimulated at an inter-stimulus interval of 40 ms to measure the paired pulse ratio (**D**) Grouped data of paired-pulse ratio of MF-CA3 EPSCs was not different between the genotypes (WT: 2.42 ± 0.15, n = 7 cells, 3 mice; A657T: 2.62 ± 0.15, n = 10 cells, 4 mice; *p* = 0.36; Mann-Whitney) (**E**) Example frequency facilitation experiment measuring MF EPSC amplitudes while elevating stimulation frequency from 0.05 Hz to 1 Hz. Inset show raw traces for each experiment in WT and A657T mice. (**F**) Grouped data for all experiments showing fold facilitation at 1Hz which was not different between genotypes (WT: 4.45 ± 1.17, n = 7 cells, 3 mice; A657T: 3.27 ± 0.38, n = 8 cells, 5 mice; *p* = 0.78; Mann-Whitney). (**G**) Timecourse for MF LTP induced by repeated tetanic stimulation (1 Hz for 1 s). Inset are traces from one experiment of EPSCs during baseline recording and 25-30 mins after induction of LTP. (**H**) Grouped data from all experiments represented as the cumulative % LTP. There was not difference in absolute LTP between genotypes (WT: 197.0 ± 38.4 %, n = 9 cells, 8 mice; A657T: 224.8 ± 42.9 %, n = 8 cells, 7 mice; *p* = 0.78; Mann-Whitney).

**Figure S2:**
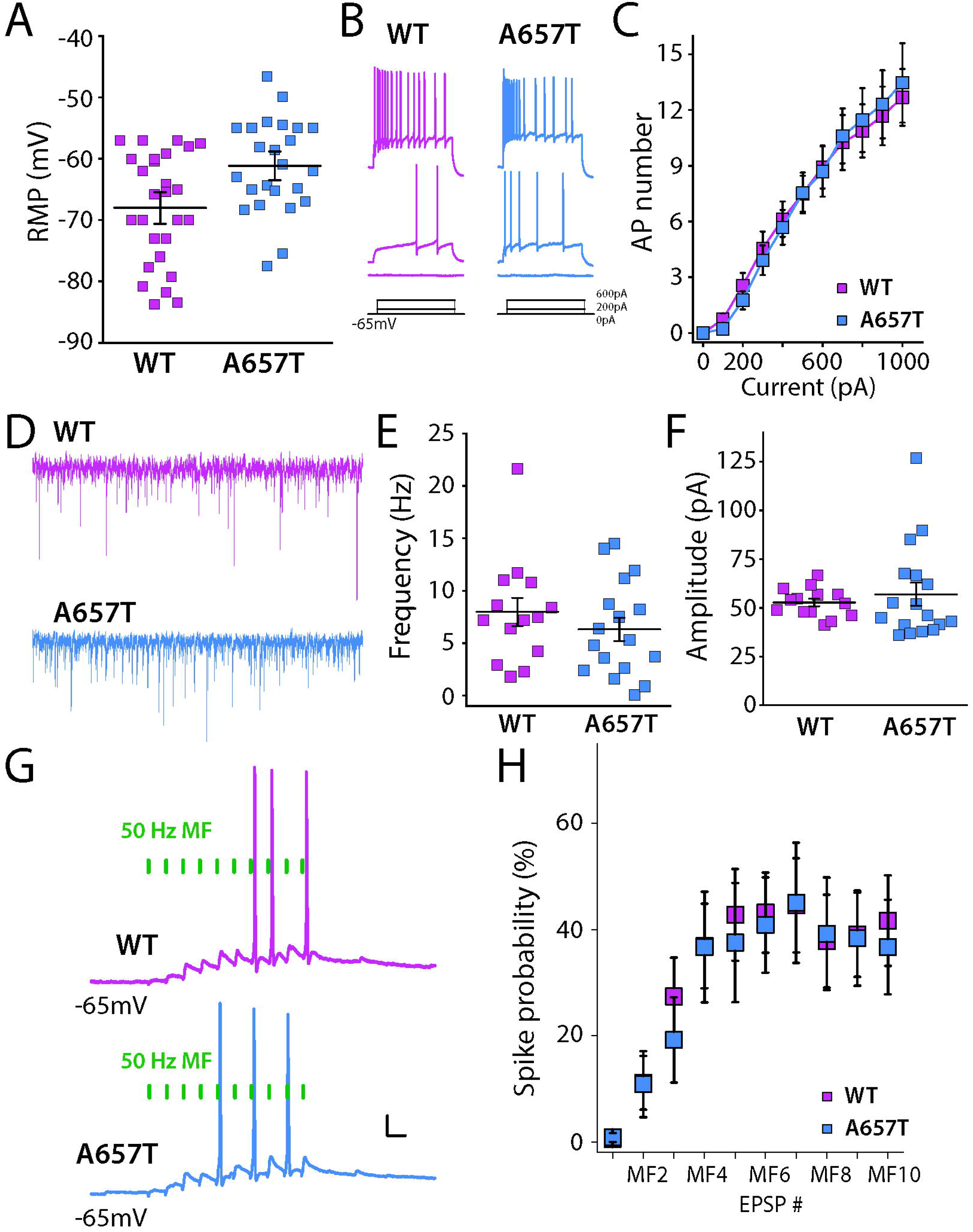
Intrinsic properties, sIPSCs, and MF-AP coupling in A657T mice. (**A**) Resting membrane potential recording in CA3 neurons is more depolarized in A657T mice than in WT mice (**B**) Example traces of APs in CA3 neurons initiated by current injection. (**C**) Full frequency input – output curve for AP firing of CA3 neurons with increasing current injection. (D) Example traces of sIPSCs recorded in CA3 neurons in WT (top) and A657T mice (lower). (E) Grouped average sIPSC frequencies for each recording (WT: 7.97 ± 1.65 Hz, n = 14 cells; A657T: 6.31 ± 1.1 Hz, n = 17 cells, *p* = 0.44, Mann-Whitney) (**F**) Average sIPSC amplitudes for each recording (WT: 52.7 ± 2.0 pA, n = 14 cells; A657T: 57.0 ± 5.9 Hz, n = 17 cells, *p* = 0.55, Mann-Whitney). (**G**) Example traces of trains MF EPSP triggered suprathreshold spikes (**H**) Grouped data of spike probability during a train of MF EPSPs in WT and A657T mice.

**Figure S3:**
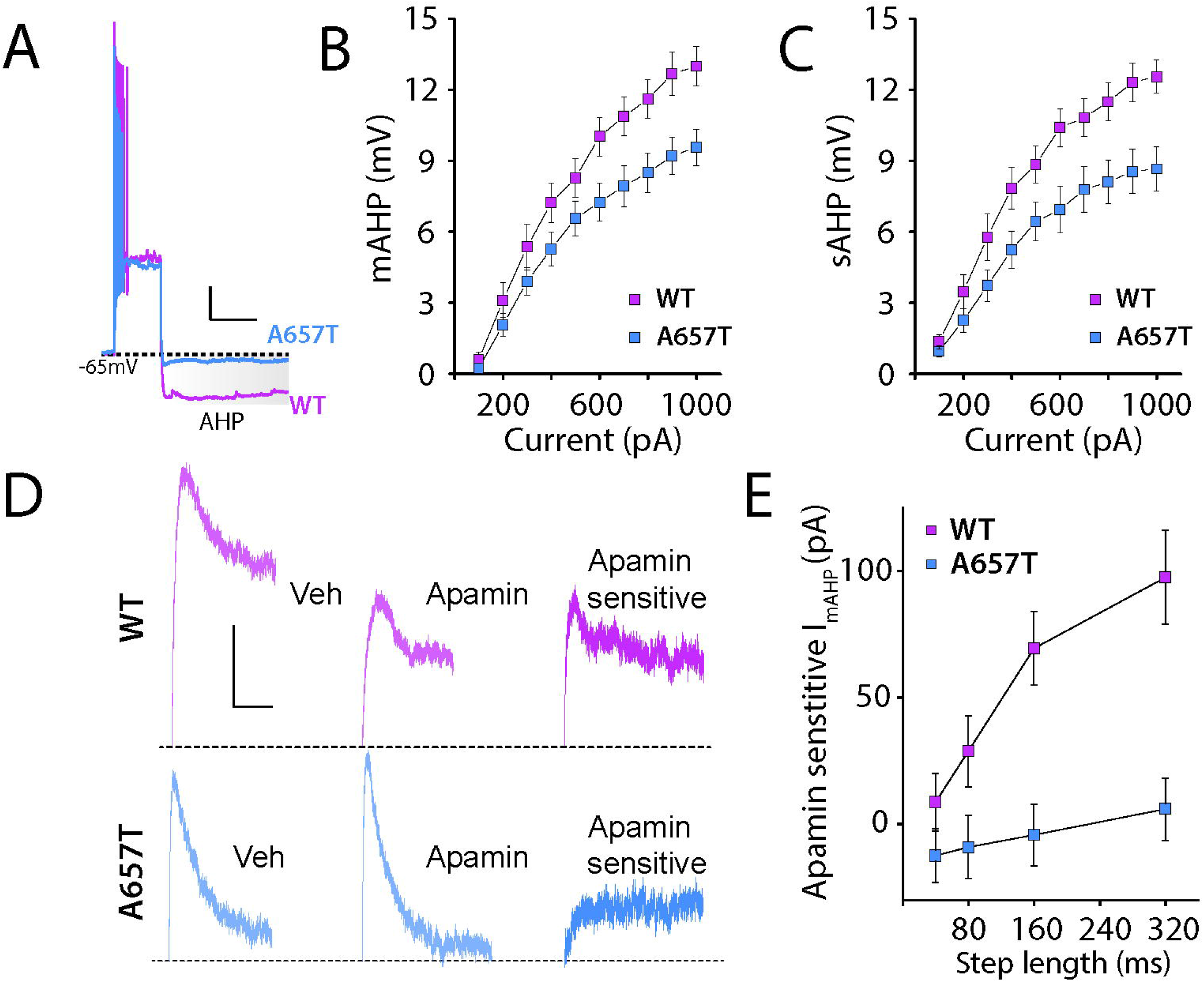
mAHP, sAHP and apamin sensitive component of I*_mAHP_* are reduced in A657T mice. **(A**) Example voltage responses to suprathreshold current steps in CA3 neurons from WT and A647T mice. The postburst AHP (highlighted) is reduced in A657T mice. Calibration 10mV, 500ms. (**B**) Input output curve for mAHP measured with increasing current injection in WT and A657T mice (**C**) measurement of sAHP with increasing current injection (**D**) Example traces of I*_mAHP_* recorded in voltage clamp before and after apamin application, and the subtracted traces (apamin sensitive). (**E**) Grouped data of the amplitude of the apamin sensitive component of the I*_mAHP_* measured with increasing current injection in WT and A657T mice.

## Notes

### Competing Interest Statement

The authors have declared no competing interest.

